# TRPM2 promotes pancreatic cancer by PKC/MAPK pathway

**DOI:** 10.1101/2020.11.09.373035

**Authors:** Rui Lin, Xunxia Bao, Hui Wang, Sibo Zhu, Zhongyan Liu, Quanning Chen, Kaixing Ai, Baomin Shi

## Abstract

**Background:** The mechanism of pancreatic cancer(PA) is not fully understanded. In our last report, TRPM2 plays a promising role in pancreatic cancer. However, the mechanism of TRPM2 is still unknown in this dismal disease. This study was designed to investigate the role and mechanism of TRPM2 in pancreatic cancer.

**Methods:** TRPM2 overexpressed and siRNA plasmid were created and transfected with pancreatic cancer cell line(BxPC-3) to construct the cell model. We employed CCK-8, Transwell, scratch wound, and nude mice tumor bearing model to investigate the role of TRPM2 in pancreatic cancer. Besides, we collected the clinical data, tumor tissue sample(TT) and para-tumor sample(TP) from the pancreatic cancer patients treated in our hospital. We analyzed the mechanism of TRPM2 in pancreatic cancer by transcriptome analysis, Westernblot, and PCR.

**Results:** Overexpressed TRPM2 could promote pancreatic cancer in proliferation, migration, and invasion ability in no matter the cell model or nude mice tumor bearing model. TRPM2 level is highly negative correlated to the overall survival and progression-free survival time in PA patients, however, it is significantly increased in PA tissue as the tumor stage increases. The transcriptome analysis, GSEA analysis, Westernblot, and PCR results indicates TRPM2 is highly correlated with PKC/MAPK pathways.

**Conclusion:** TRPM2 could directly activate PKCα by calcium or indirectly activate PKCε and PKCδ by increased DAG in PC, which promote PC by downstream MAPK/MEK pathway activation.

---

Pancreatic cancer is one of the most devastating cancer, of which median survival is 3-5 months and the five year survival rate is less than 4%^[1]^. According to the statistics, the incidence and mortality rate gradually increased by 1.3% and 1.25% per year in China in the period of 2004-2015^[1]^. The real incidence rate must be higher than the statistics because the statistics data^[1]^ covered only 12% of the nation’s population^[2]^. Despite the advance in surgical techniques and adjuvant therapy, the prognosis doesn’t have changed much in over four decades. The recent median overall survival have only improved by 5 months on the basis of the new combined therapy for metastasis pancreatic cancer^[3]^. The molecular mechanism of pancreatic cancer is desperately investigated worldwide nowadays for a better understanding of this dismal disease. In last report by our research^[4]^, TRPM2 played a promising role in promoting pancreatic cancer. Here, we designed new experiments to investigate the role and mechanism of TRPM2 in pancreatic cancer.

## Methods

### Collection of clinical data and specimens

Collected clinical data and pathological specimens of tumor tissue (TT) and paratumor tissue (TP) from 64 patients with pancreatic cancer who were treated in Tongji Hospital Affiliated to Tongji University from May 2017 to May 2019. The patients’ tumor stage were difining according to the pathological results and AJCC TNM staging. All patients were followed up. The end date of follow-up was December 31, 2019. Transcriptome sequencing was performed on the surgical specimens of 12 patients, of which three in each stage of pancreatic cancer. TRPM2 immunofluorescence staining was performed on TT and TP specimens. All patients signed informed consent.

### Western-blot

Part of the tissues of TT and TP in PDAC patients was homogenized in lysis buffer containing protease and phosphatase inhibitor (Sigma-Aldrich).Protein samples were separated on SDS-PAGE gel and transferred to nitrocellulose membrane. After blocking, incubate with Ras, Raf-1, PSPH, OASL, PKC, METTL3, HIST1H2AE, cPLA and AQR antibodies (abcam) at 4°C overnight. After incubating with the secondary antibody at 20°C for 1 hour, the membrane was washed 3 times. The immune response zone is detected by the ECL detection system. The protein bands were quantified using Image J software (NIH, Bethesda, MD, USA).

### Lentivirus transfection to construct TRPM2 overexpressing/interfering BXPC-3 cells

The pancreatic cancer cell line BXPC-3 cells (CinoAsia Co., Ltd.) were inoculated into a 24-well plate (1×10^5^/well), and the added medium ((RPMI 1640; 10% fetal bovine serum, Gibco; Thermo Fisher Scientific, Inc., Waltham, MA, USA)) with a volume of 500ul. Incubate at 37°C, 5% CO2. The number of cells during lentivirus transfection is about 2×10^5^/well. The next day, replace the original medium with 0.5ml fresh medium containing 10μg/ml polybrene, and add 10ul of 1×10^8^TU/ml TRPM2 overexpression/interference lentivirus suspension (Shanghai GenePharma Co., Ltd), After incubating at 37°C for 24h, change the medium and replace the virus-containing medium with 1ml of fresh medium. After 72 hours of infection, observe the expression of GFP with a fluorescence microscope, and add medium containing the optimal selection concentration of antibiotics to screen the cells (about a week). The screened cells are further purified using the limiting dilution method, and the cultures are expanded after purification. After the cells have grown to a sufficient number, collect the cells and extract RNA and perform RT-PCR verification. The cells that are verified as positive can be used for subsequent experiments or frozen storage.

### RNA extraction and RT-PCR

Total RNA was extracted from cells and tissues using Trizol reagent, and the quality of extracted RNA was confirmed using NanoDrop 1000. After reverse transcription, a PCR reaction system was prepared, and qPCR analysis was performed by real-time detection system through SYBR green I dye (Takara) detection. According to the PCR standard reaction curve, the original Ct values of the target genes TRPM2, Ras, Raf-1, PSPH, OASL, PKC, METTL3, HIST1H2AE, cPLA and AQR were obtained, and the 2^-ΔΔCt^ method was used for semi-quantitative analysis.

Table 1 showed the detailed primer sequence of each gene.

**Table 1.**
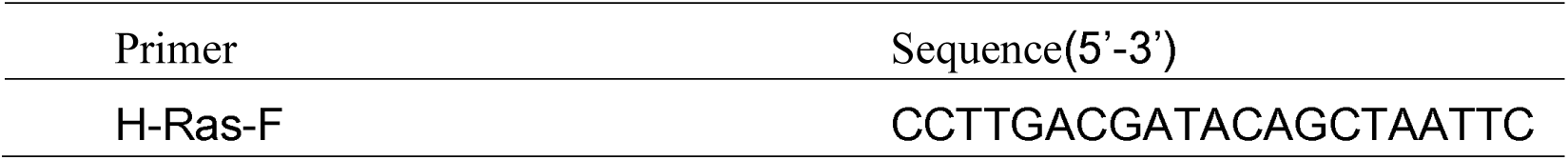
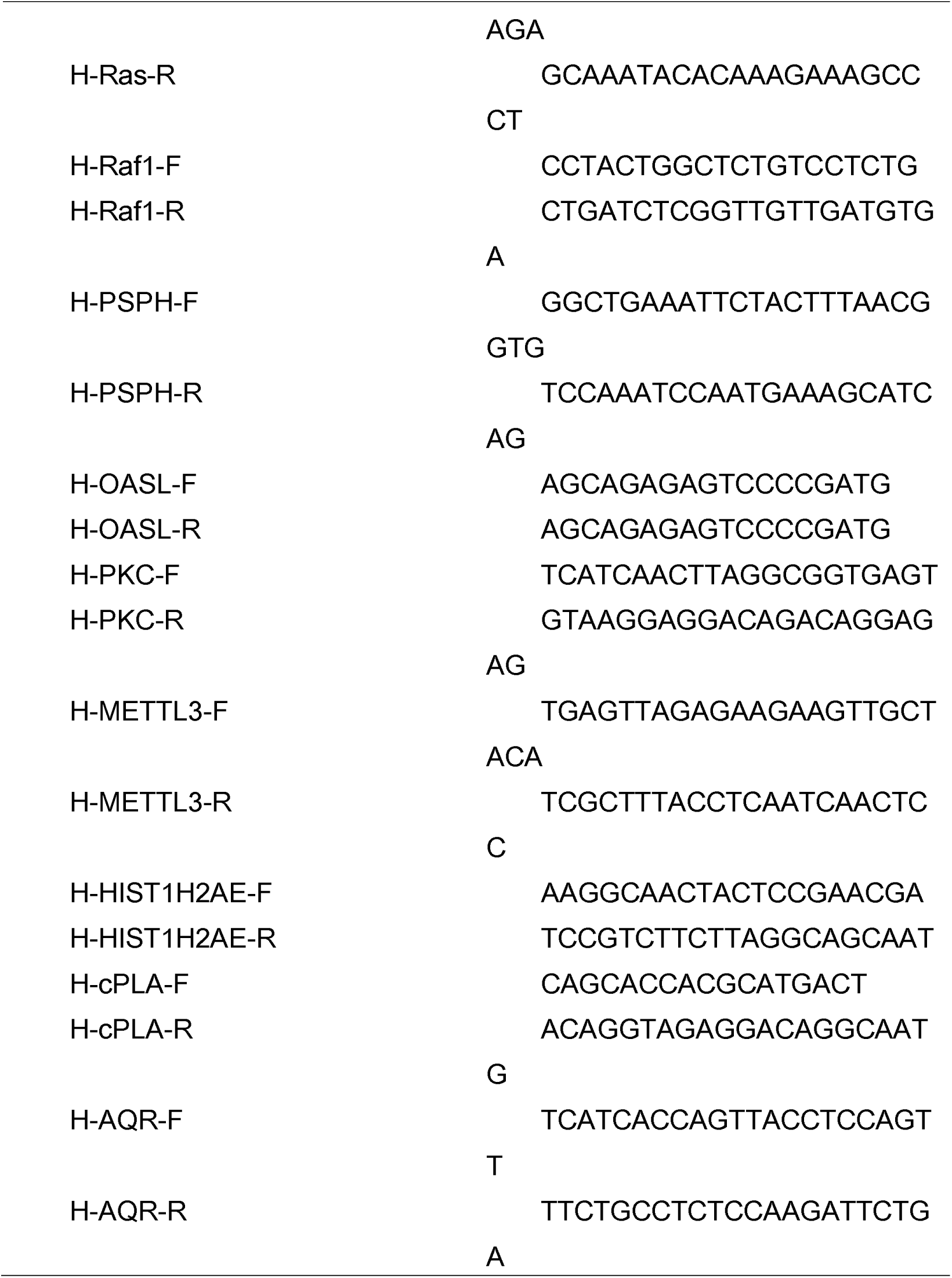
Primer sequence

### CCK-8 assay

The constructed NC, TRPM2 vector, empty vector, TRPM2 siRNA, and Scramble siRNA BXPC-3 cells were respectively seeded in 96-well culture plates (1×10^4^/well) and placed at 37°C, Incubate overnight in a 5% CO_2_ incubator.Wash the cells inoculated the previous day with PBS twice, then add 100ul 1640 medium for use. Discard the old solution at 0h, 24h and 48h after changing the medium, add 95ul of medium and 5ul of CCK-8 to each well of the 96-well plate to be tested, and incubate at 37□ for 2h. A microplate reader (BioTek Instruments, Inc., Winooski, VT, USA) was used to measure the OD value of each experimental well at 450 nm, and to detect changes in cell proliferation ability of each group.

### Transwell assay

Inoculate the constructed NC, TRPM2 vector, empty vector, TRPM2 siRNA, and Scramble siRNA group cells into the wells of Matrigel plate containing serum-free RPMI 1640 medium (5×10^4^cells/well). The lower well contains 500 µl of complete medium (RPMI 1640 and 10% fetal bovine serum). After incubation at 37°C for 48 hours, gently remove cells that have not migrated through the well with a cotton swab. The cells in the lower chamber were fixed with 5% glutaraldehyde for 10 minutes and stained with 1% crystal violet in 2% ethanol at room temperature for 20 minutes, photographed and counted.

### Wound healing assay

The constructed NC, TRPM2 vector, empty vector, TRPM2 siRNA, and Scramble siRNA group cells were inoculated into a 24-well culture plate (1×10^5^/well), and placed at 37°C, 5% CO_2_ for 24h. Remove the medium and scratch the surface of the inoculated cells with a 10ul pipette tip and mark it. Wash gently twice with PBS. Add 1 ml RPMI 1640 medium. Photograph the scratches at 0h and 24h.

### Construction of nude mice bearing tumor model

Cells in the logarithmic growth phase were digested with 0.25% trypsin, centrifuged at 800 rpm for 3 min, and the supernatant was discarded to collect the cells. Count the cells after resuspending them in PBS, and adjust the cell concentration to about 6×10^7^/ml, put on ice for use.

Prepare 9 male BALB/c nude mice (purchased from Shanghai Experimental Animal Center, Chinese Academy of Sciences) of 4-6 weeks of age with no specific pathogens with average weight of 17±2g. We divided the mice into three groups, three in each group, and inoculated them with BXPC-3 cells in overexpression(OE), normal control(NC) and siRNA interfering(siRNA) groups. We aspirated the cell suspension with a 1ml syringe (shake the cell suspension before mixing), and slowly injected 150ul suspension into the subcutaneous area of axilla of nude mice. After the tumor cells were inoculated, the general condition and growth of the nude mice were closely observed. The nude mice were kept in a SPF air clean laminar flow rack at a constant temperature of 22-28°C and a constant humidity of 45-60%. Measure tumor volume once a week. After 6 weeks, the nude mice were euthanized. The tumors were carefully peeled off and removed. The tumors were placed neatly and photographed.

The tumor volume was measured with a ruler and weighed on the scales. Volume formula: volume = long diameter×short diameter^2^×1/2. The study was approved by the ethics committee of Tongji Hospital affiliated to Tongji University.

### Immunofluorescence

The TT and TP tissues of PDAC patients and nude mice were fixed and sectioned. TRPM2 antibody (abcam) was used for immunofluorescence detection to analyze the difference in expression of TRPM2. The slices were scanned for image acquisition and image J was used to analyze the optical density value.

### Transcriptome sequencing analysis

RNA was extracted from TT and TP tissues of PDAC patients and nude mice, and reverse transcribed into cDNA for PCR amplification. The PCR amplification reaction product was purified and subjected to quality control in a 96-well plate. After dilution, the samples were sequenced using Hiseq2500 sequencer (Illumina, USA). Trimmomatic processed the raw data and used featureCounts software to quantify the expression level of each gene. Perform principal component analysis (PCA) and cluster analysis (HCA) on the samples. And use R package “ComplexHeatmap” and “pheatmap” to draw a heatmap, “stat” package to analyze differentially expressed genes (DEG), with P<0.05 as the standard. RNAseq deconvolution to decipher tumor microenvironment by R package ‘immunedeconv’. Use the ‘Venn Diagram’ online tool to generate DEG intersections, and use the “KEGG profile” for path analysis. DAVIDv6.7 performed gene ontology analysis (GO). Gene Set Enrichment Analysis (GSEA) human mouse pancreatic cancer signal pathway intersection and related genes.

## Results

### Clinical data indicated that high expression levels of TRPM2 in PDAC patients are associated with poor prognosis

A total of 64 patients with pancreatic cancer were enrolled in this research project, including 38 men and 26 women, whose average age was 63.1±7.4 years. There were 37 patients’ tumors located in pancreatic head, 12 located in uncinate process of pancreas, and 15 located in the body and tail of pancreas. All patients received surgery in our hospital, in which 39 received pancreaticoduodenectomy, 4 received pancreatectomy of middle section, 8 received tail pancreatectomy, and 13 received palliative surgery. All specimens were diagnosed as pancreatic ductal adenocarcinoma by pathologist. According to AJCC TNM stage, the patients were subdivided into 4 different stage groups depending on their pathology results. There were 6 patients with stage I (including IA and IB), 22 patients with stage II (including IIA and IIB), 27 patients with stage III, and 9 patients with stage IV. Table 2 showed the patient baseline characteristics. There were no significant difference for sex(*P*=0.94), age(*P*=0.57), and ECOG performance score(*P*=0.118) among four groups. However, the tumor grade, median survival, and median progression-free survival were statistically significant different among four groups(*P*<0.05).

**Table 2.**
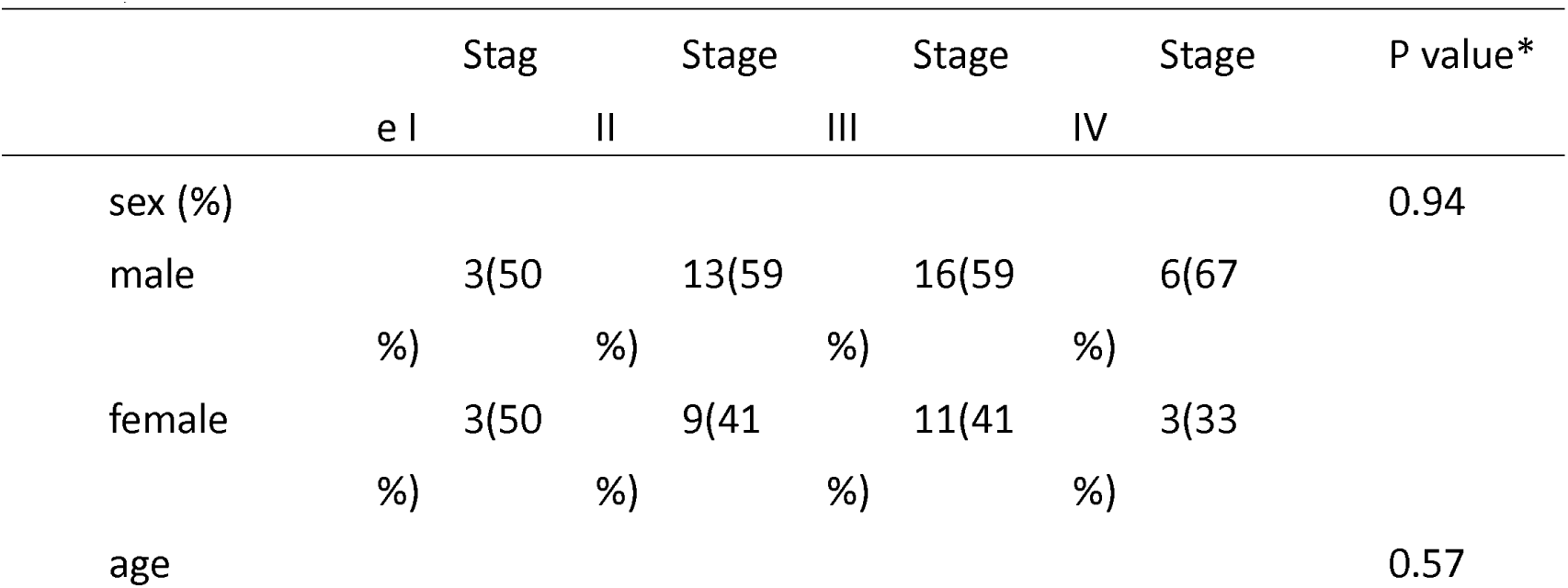
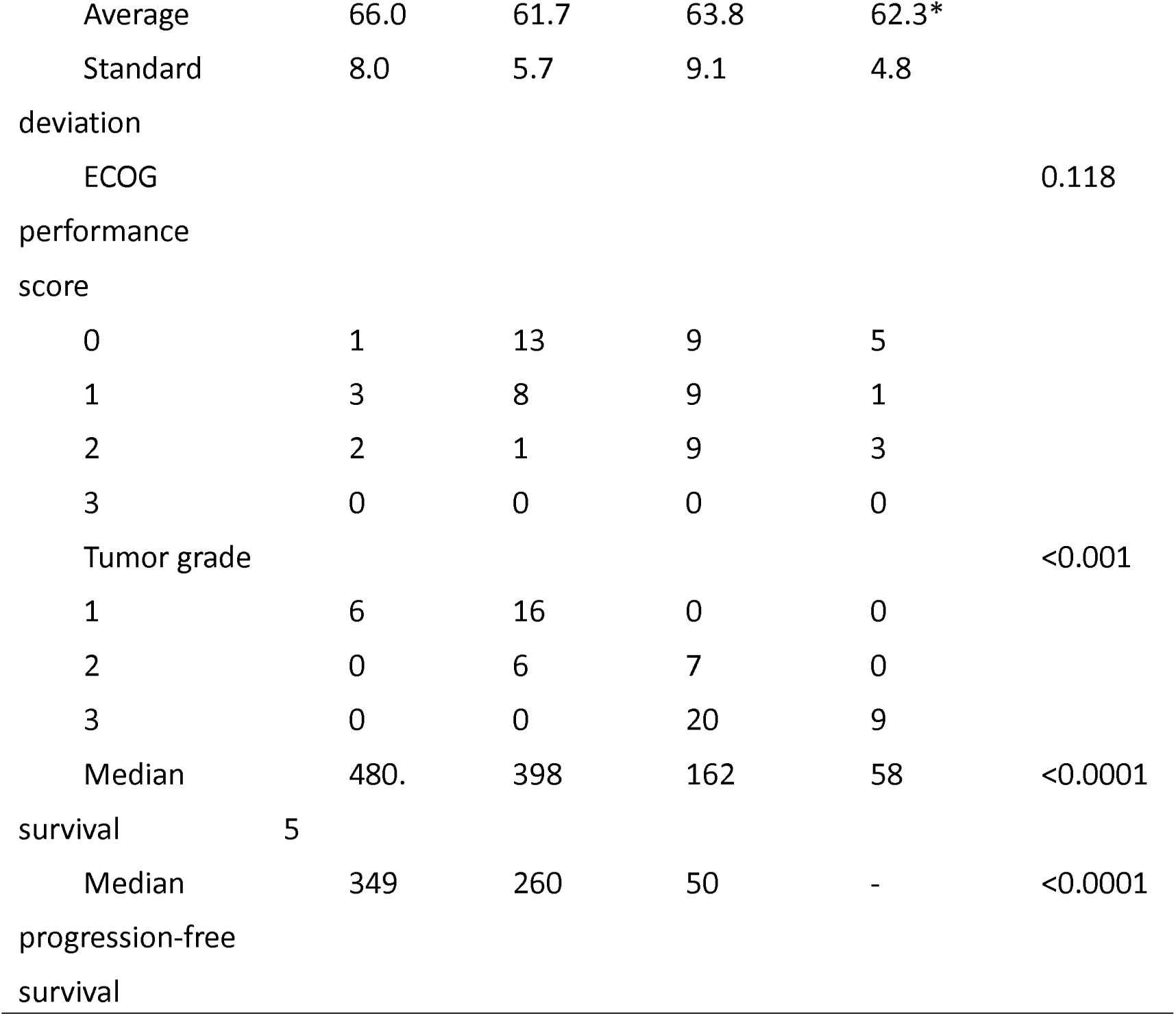
Patient baseline characteristics (one-way ANOVA, chi-square test, * P <0.05)

The follow-up data showed that the patient’s progression-free survival (Figure 1a) and overall survival (Figure 1b) decreased significantly as the tumor stage increased. After performing RNA sequencing for each tumor and para-tumor specimen, the sequencing data was in good quality control(Figure 1c, d, e). We found that TRPM2 was the only molecule of TRPM family that highly expressed in tumor than para-tumor specimens in the RNA level(*P* <0.05, Figure 2a). Immnunostaining images for different stage groups showed TRPM2 expresseion in protein level was statistically significantly increasing as the tumor stage increased and tumor specimens had higher level than para-tumor specimens(*P*<0.05, Figure 2b). The OD value of TRPM2 expression in immunostaining images were used in calculation of Pearson coefficient. The Pearson coefficient between TRPM2 level and survival was -0.88, TRPM2 level and recurrence(progression-free survival) was -0.85.(Figure 2c). However, the Pearson coefficient between TRPM2 and tumor stage was 0.81(Figure 2c).

**Figure 1.**
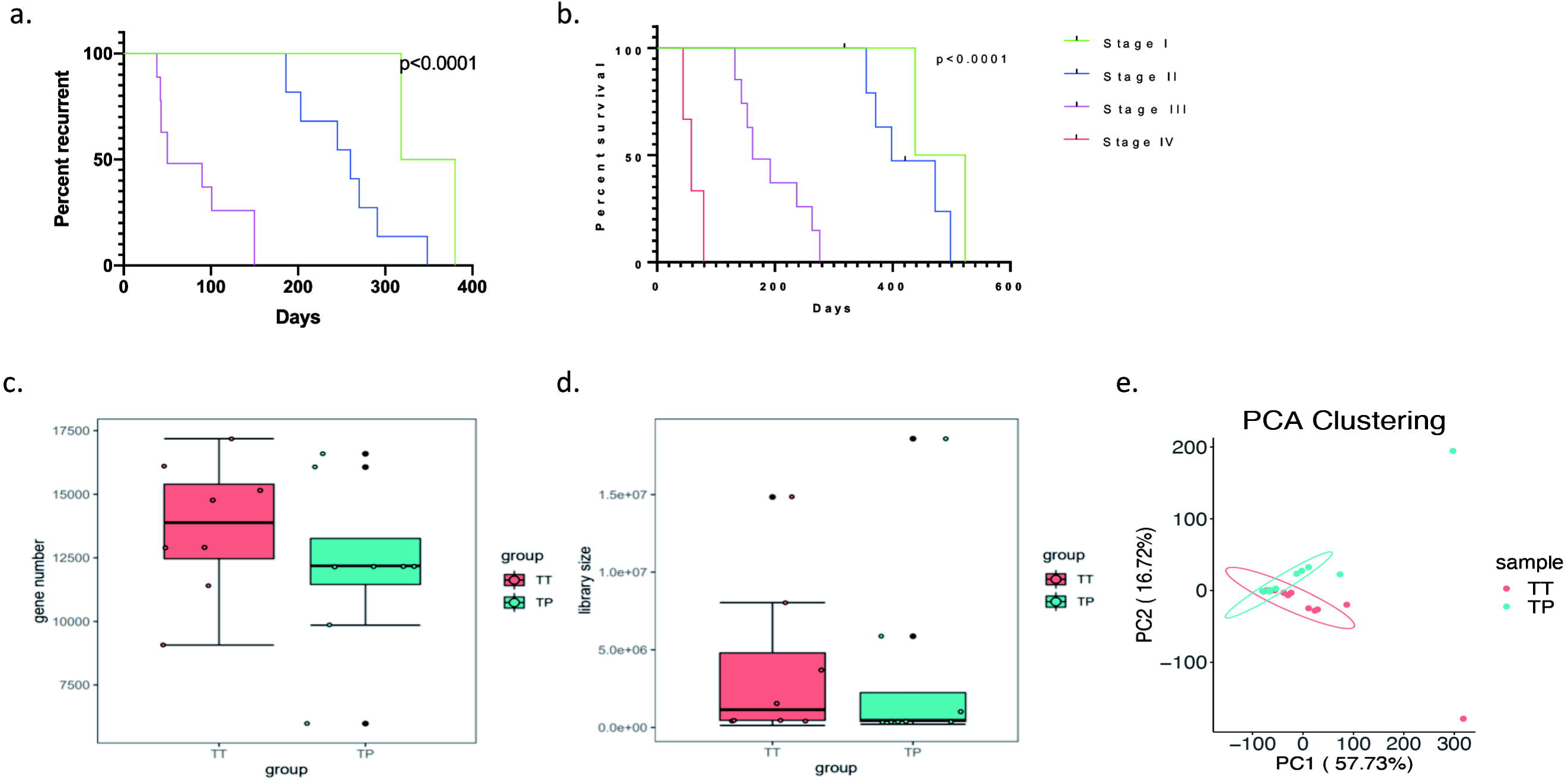
(a) (b) The Kaplan-Meier plots showed that the progression-free survival and overall survival of patients decreased significantly as the patients’ tumor stage increased. The significance was tested by Log-rank test; (c) Quality Control for patients’ tumor(TT) or para-tumor(PT) tissue: The plot showed no significant different gene number sequenced in two groups; (d) Quality Control for patients’ tumor(TT) or para-tumor(PT) tissues: Library size sequenced showed no significant difference between two groups; (e) Quality Control for patients’ tumor(TT) or para-tumor(PT) tissue: PCA clustering showed that there were significant differences between patients’ TT and TP tissues.

**Figure 2.**
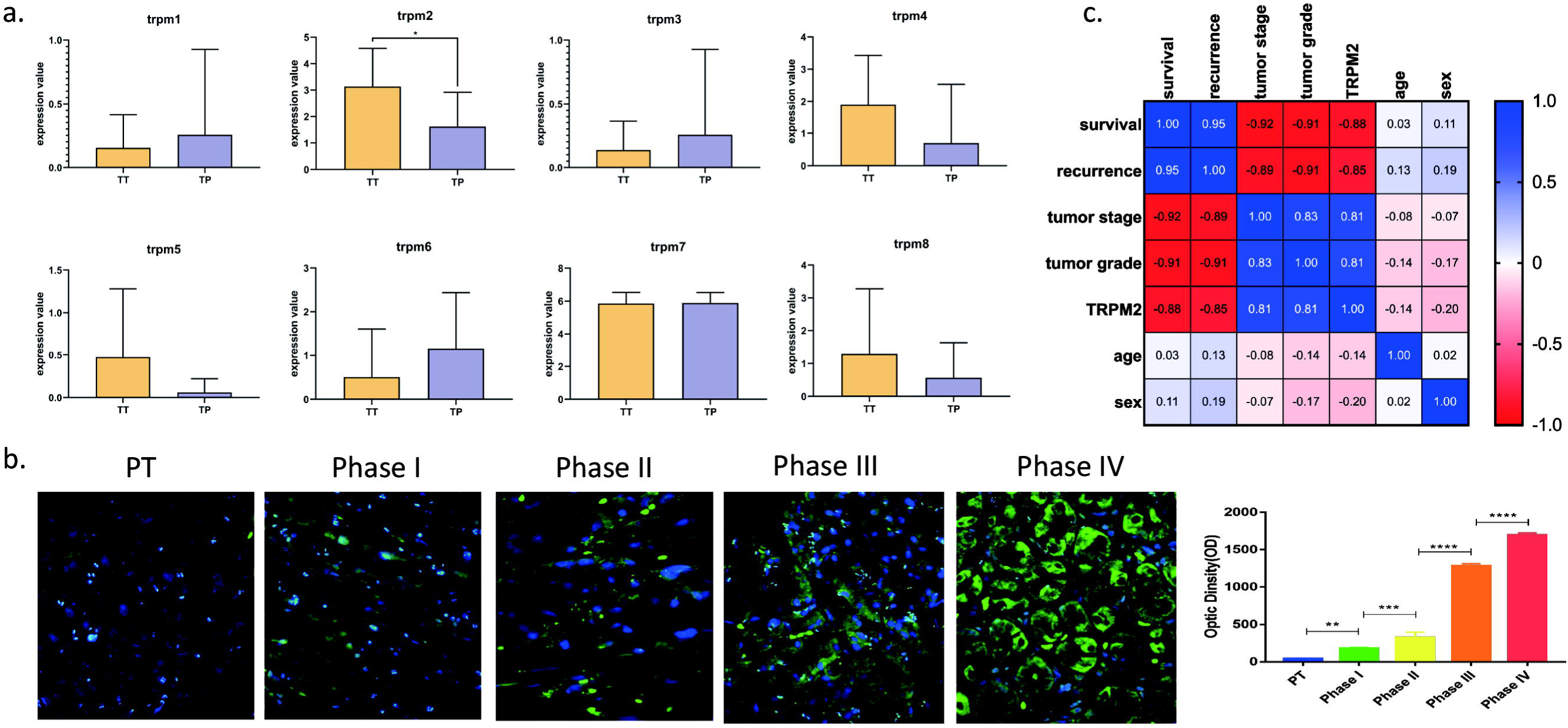
(a) The mRNA level of TRPM 1-8 in transcriptome sequencing data for TT and TP in PC patients(* *P*<0.05); (b) Immunofluorescence staining of TRPM2 in PT and different stages of TT tissues(Stage I to IV, ** *P*<0.01, *** *P*<0.001, **** *P*<0.0001); (c) Pearson’s coefficient in Multivariate Analysis for patients’ clinical and sequencing data.

### TRPM2 promotes the proliferation, migration and invasion of BxPC-3 pancreatic cancer cells

After BxPC-3 cells were transfected with lentivirus, the relative mRNA expression level of TRPM2 in NC, empty vector, Scramble siRNA, TRPM2 siRNA, and TRPM2 vector groups were 1±0.08, 1±0.1, 0.9±0.11, 0.03±0.01, 4.53±0.23. The results showed that BXPC-3 cell models with overexpressed and interfered TRPM2 were successfully constructed. The expression level of TRPM2 was significantly higher in TRPM2 overexpressed group than any other group (*P*<0.0001) and that in TRPM2 siRNA group was significantly lower than any other group(*P*<0.0001Figure 3a).

**Figure 3.**
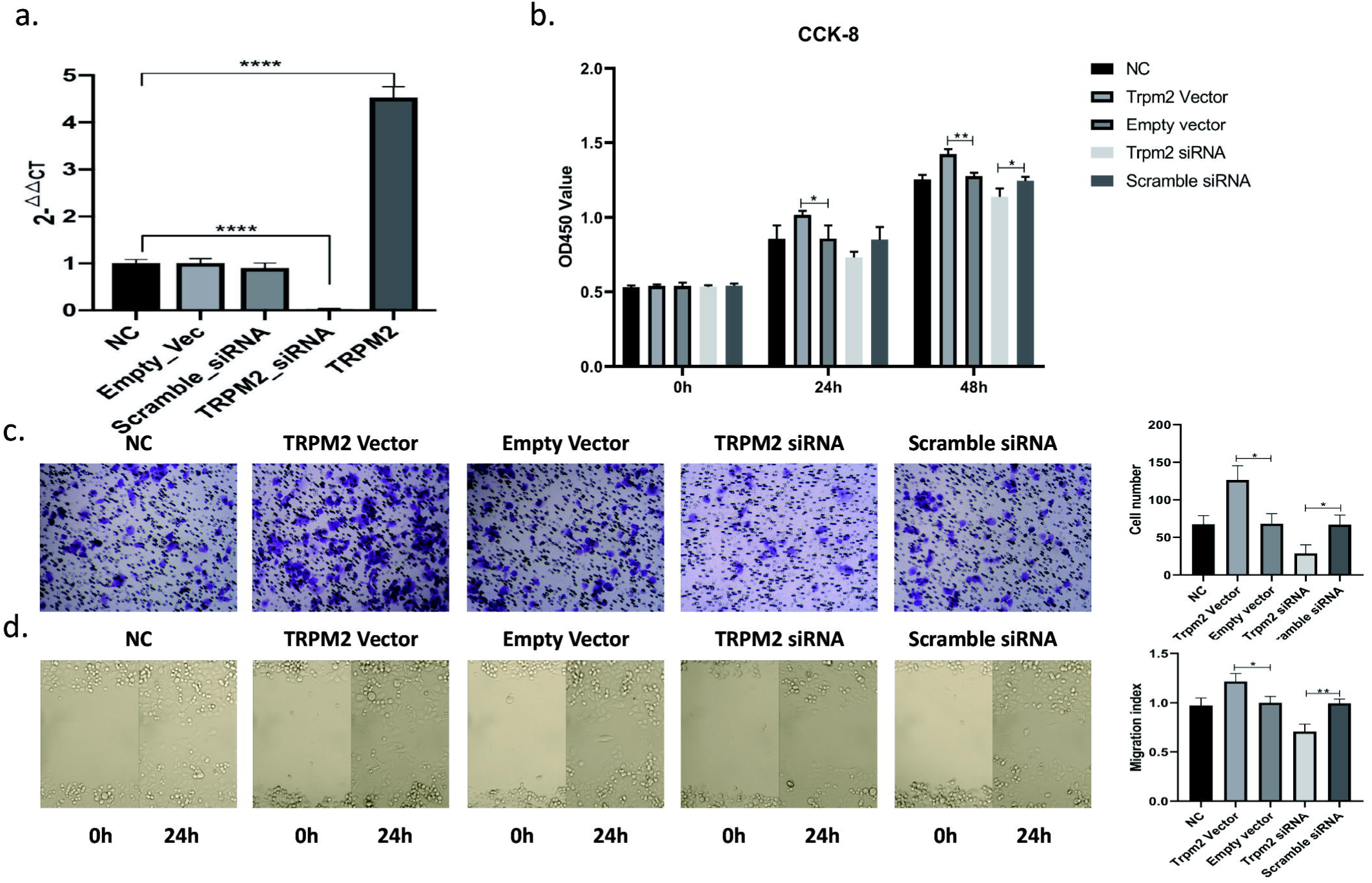
(a) RT-PCR results in different experimental groups of BxPC-3 cells: TRPM2 overexpressed(TRPM2), TRPM2 siRNA, scramble siRNA, empty vector(Empty_Vec), and normal control(NC) groups. **** *P*<0.0001; (b) CCK8 assay results of different groups at 0-, 24-, and 48-h time intervals. * *P*<0.05, ** *P*<0.01, siRNA, small interfering RNA; CCK8, cell counting kit-8; OD, optical density. (c) Transwell assay results showed that the TRPM2 overexpression group has a stronger invasive ability and TRPM2 siRNA group has a weaker invasive ability. Images under microscopy(magnification x100). * *P*<0.05 (d) Scratch wound healing assay results. Images of all groups at 0- and 24-h time intervals post injury(magnification x100). * *P*<0.05, ** *P*<0.01

In CCK-8 assays, twenty-four hours after cell culture for different groups, the OD value of the TRPM2 overexpression group increased significantly compared with other groups (*P*<0.05). After forty-eight hours, the OD value of the TRPM2 overexpression group increased significantly compared with other groups (*P* <0.01), and the OD value of the TRPM2 siRNA group decreased significantly (*P* <0.05). This indicates that TRPM2 has a proliferation-promoting effect on BxPC-3 pancreatic cancer cells (Figure 3b).

In the Transwell assay, after 48 hours of cell inoculation, an average cell count of 5 fields was randomly taken under the X100 microscope. The cell count of TRPM2 overexpression group was significantly increased compared with other groups (*P*<0.05), while the cell count of TRPM2 siRNA group was significantly decreased compared with other groups (*P*<0.05). It showed that TRPM2 increased the invasiveness of BxPC-3 pancreatic cancer cells (Figure 3c).

In the scratch wound healing assay, the wound healing level was calculated at two time points, 0 and 24h. As shown in Figure 3d, the cell migration ability was significantly enhanced in TRPM2 overexpressed group than that of the other groups (*P*<0.05). However, the cell migration ability in TRPM2 siRNA group was significantly weaker than that of the other groups (*P*<0.01). It showed that TRPM2 could promote migration of BxPC-3 pancreatic cancer cells.

### TRPM2 function experiment of nude mice tumor bearing model

After six weeks implantation, the tumors were dissected from the nude mice to measure the volume and weight.(Figure 4a). The volume of the tumors were significantly higher in TRPM2 overexpressed(OE) group than in NC group and TRPM2 siRNA group in each week from the second week to the sixth week after implantation(*P*<0.0001, Figure 4b). The weight of the tumors were significantly higher in OE group than in NC(*P*<0.05) and siRNA group(*P*<0.001) in the sixth week after implantation(Figure 4c). It suggested that TRPM2 has the effect of promoting the growth of implanted tumors in nude mice.

**Figure 4.**
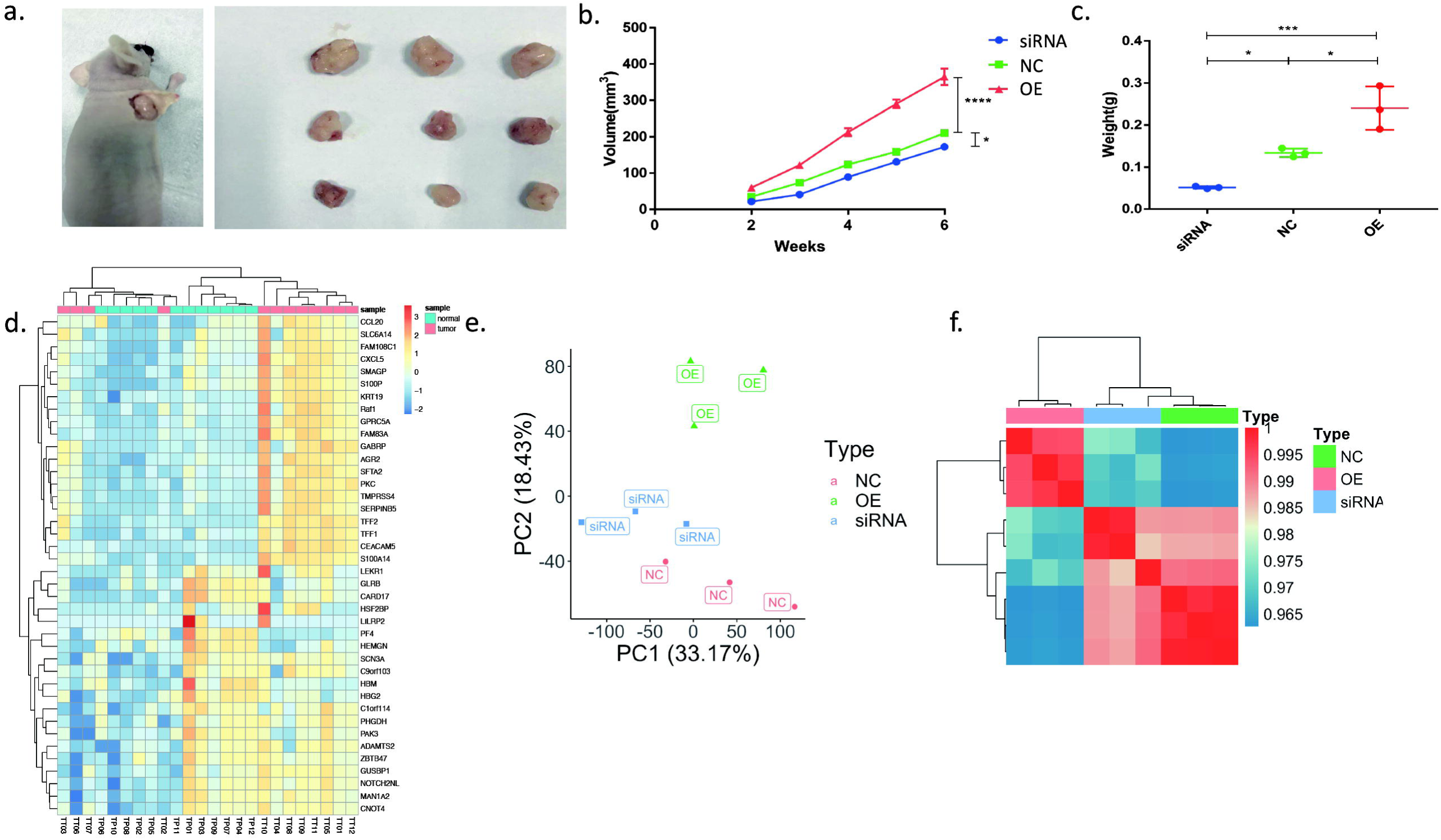
(a) Tumors of nude mice; (b)Weekly volume change of nude mice implanted tumors (c) Weight change of nude mice implanted tumors after six weeks. (OE: TRPM2 overexpression, NC: normal control, TRPM2 siRNA: TRPM2 interference expression group, * *P*<0.05, *** *P*<0.001, **** *P*<0.0001); (d) Heatmap of top 40 differentially expressed genes(DEG) between TT and TP groups of patients’ PC samples. (e) PCA clustering showed significant differences among NC, OE, and siRNA groups. (f) Heatmap of DEGs among NC, OE, and siRNA groups.

### TRPM2 could promote pancreatic cancer through PKC/MEK pathway

We sequenced the transcriptome data in both human tissues(TT vs. TP) and mice implanting tumors(siRNA vs. NC vs. OE). In PCA analysis, we could find that there were significant differences between TT and TP group (Figure 1e). The top 20 differentially expressed genes(DEGs) between TT and TP groups were displayed in the heatmap (Figure 4d). The similar outcomes were also seen in tumor-bearing nude mice models(Figure 4e, f).

The DEGs between TT vs. TP groups could enrich in 167 different pathways in the GSEA database(Figure 5b). Similar outcomes were found in DEGs between OE vs. siRNA groups in nude mice, in which 169 different pathways were enriched in the GSEA database(Figure 5b). There were 18 common pathways between human and mice samples(Figure 5b).

**Figure 5.**
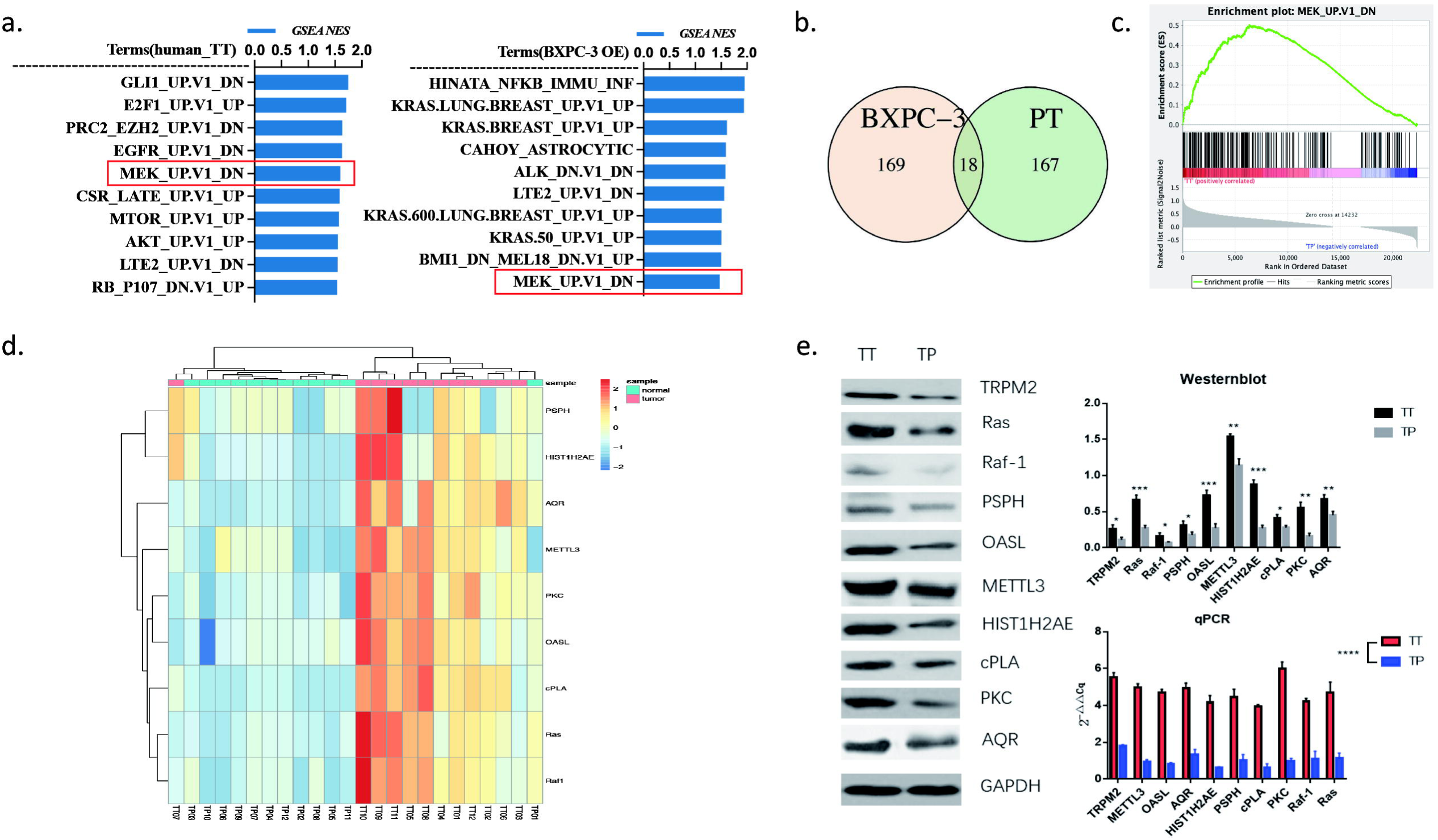
(a) Top ten standard NES of TT vs. TP in patients and OE vs. NC in nude mice; (b) Intersection of signal pathways enriched in tumors of nude mice and human; (c) GSEA analysis of enrichment plot of MEK pathway; (d) Screening of key genes of the MEK pathway signaling pathway in TT vs. TP groups; (e) Using Westernblot and qPCR to verify the expression of MAPK/MEK key molecules in patient’s TT and TP groups (\**P*< 0.05, ** *P*<0.01, *** *P*<0.001, **** *P*<0.0001).

The top 10 pathways from 18 common pathways between human and mice samples were listed in Figure 5a. As shown in the figure, MEK pathways ranked the top 10 in both human and mice samples(Figure 5a, c).

The downstream pathways of TRPM2 from KEGG database was shown in Figure 7. The TRPM2 may promote pancreatic cancer through PKC/MEK pathways.

**Figure 6.**
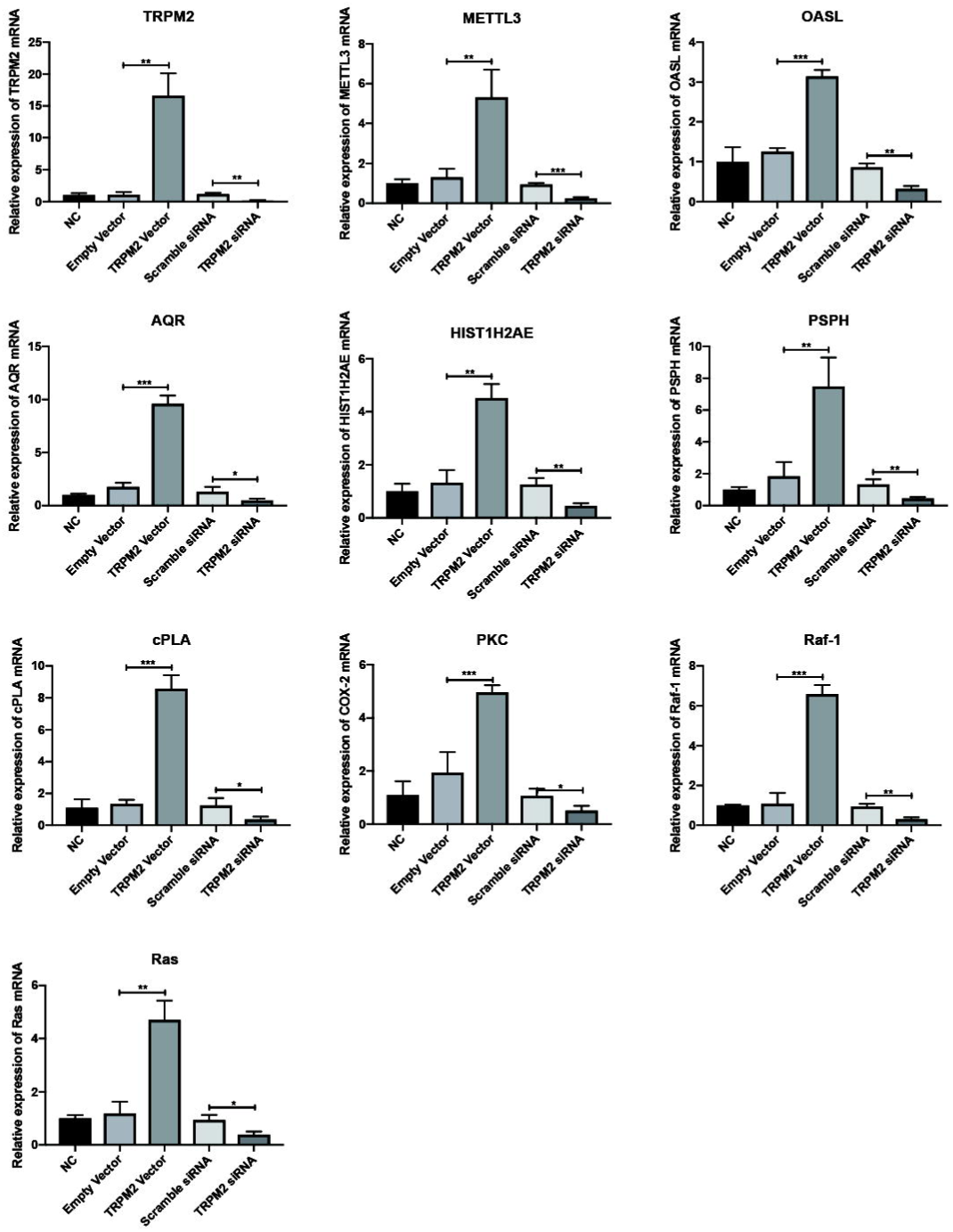
Expression of MAPK/MEK pathway key molecules in BxPC-3 stable transfection groups. NC, normal control; Empty Vector, BxPC-3 without any plasmid transfection; TRPM2 vector, TRPM2 overexpression group; scramble siRNA, BxPC-3 cells with scramble sequence of small interfering RNA transfection group; TRPM2 siRNA, BxPC-3 cells with TRPM2 small interfering RNA transfection group. ** *P*<0.01, *** *P*<0.001.

**Figure 7.**
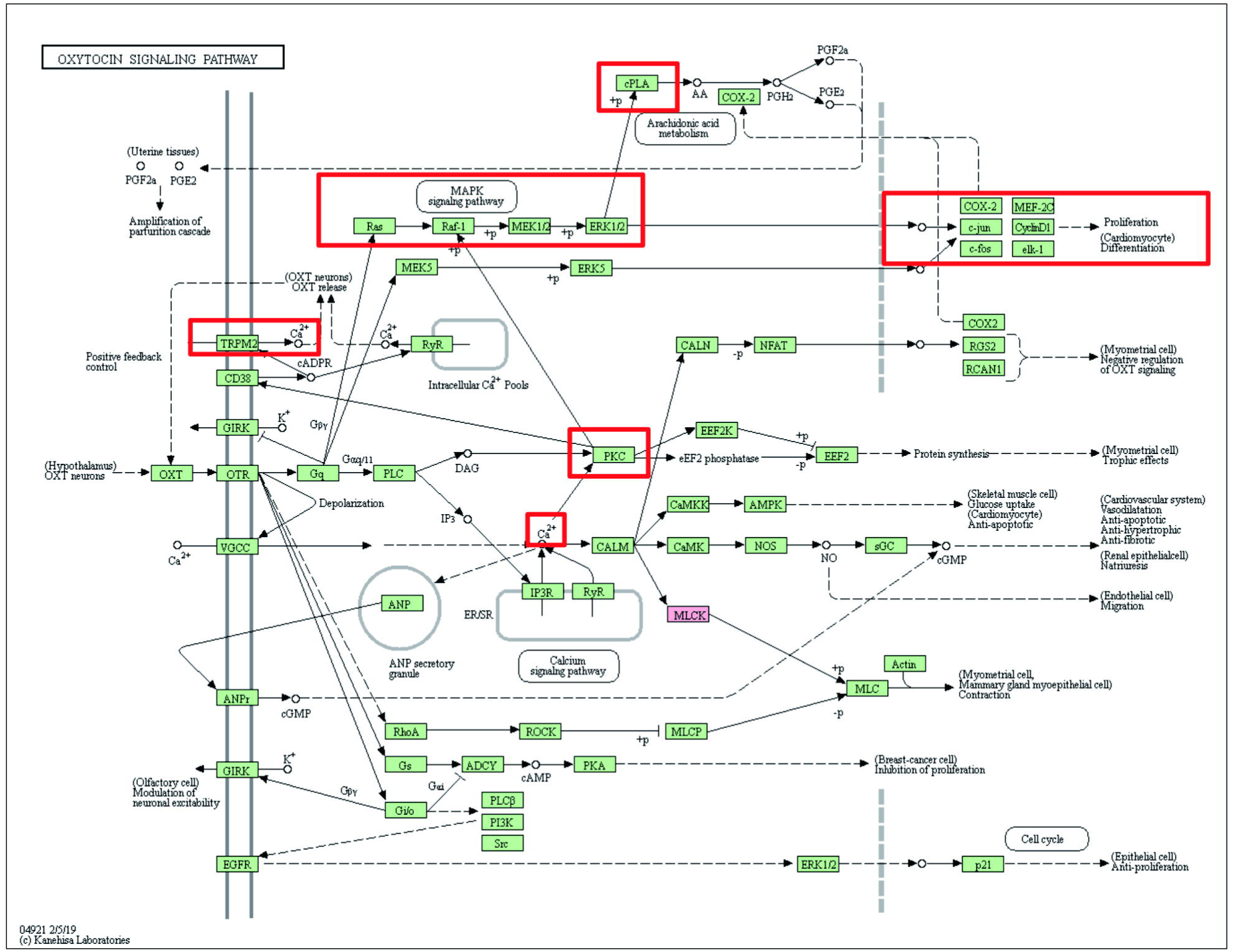
Downstream pathways of TRPM2 in KEGG database. The red-square marking pathway coincides with the pathway we found in this article. TRPM2 could increase calcium inflow to activate PKC. MAPK pathway is the downstream pathway of PKC which could promote cell proliferation.

For identifying the mechanism of TRPM2/PKC/MEK, we picked out nine DEGs, including Ras, Raf-1, PSPH, OASL, METTL3, HIST1H2AE, cPLA, PKC, and AQR, in the PKC/MEK pathways from human sequencing data and drew the heatmap(Figure 5d). In the heatmap, these 9 genes were expressed significantly higher in the TT than TP tissues. There were significant differences between TT and TP tissues in each of the nine genes no matter in protein(Westernblot) or mRNA(qPCR) level(Figure 5e). The similar results were also found in BxPC-3 models. Each of the 9 genes plus TRPM2 were significantly higher expressing in TRPM2 overexpressed group than NC, empty vector, and scramble siRNA groups(Figure 6). Each of the 9 genes plus TRPM2 were significantly lower expressing in TRPM2 siRNA group than NC, empty vector, and scramble siRNA groups(Figure 6).

The above results indicate that TRPM2 was closely related to its downstream regulatory pathway PKC/MEK in promoting the proliferation and invasiveness of human pancreatic cancer cells°

## Discussion

Transient receptor potential(TRP) channels are a cluster of ion channels which have many physiological functions^[5]^. The subfamily of TRPM(Melastatin) participate in modulation of cell proliferation and survival^[6]^. One of these is TRPM2, which is the second member of the TRPM subfamily to be identified. Human TRPM2 is permeable to Ca^2+^, Na^+^, and K^+^ which consists of 1503-amino acid^[7]^.

TRPM2 is highly expressed in many kinds of cancer, like gastric cancer^[8]^, breast cancer^[9]^, prostate cancer^[10]^, and lung cancer^[11]^. It was reported that TRPM2 could promote cancer proliferation through regulation of oxidative stress^[12]^, autophagy^[8]^, and mitochondrial function^[13]^.

In 2018, our group first report the role of TRPM2 in pancreatic cancer^[4]^. Plus the results in this article, we have demonstrated that TRPM2 could promote proliferation and invasion ability in vitro in both PANC-1 and BxPC-3(Fig. 3) pancreatic cancer cell lines. The nude mice tumor bearing model results in this article is another evidence that support TRPM2’s promotion effect in pancreatic cancer(Fig. 4a, b, c).

We found that TRPM2 was significantly higher expressed in human pancreatic cancer tissue than in para-tumor tissue in both mRNA level(Fig. 2a) and protein level(Fig. 2b). The level of TRPM2 also increased as the patients’ TNM stage increased(Fig. 2b), meanwhile, the patients’ overall survival(OS) period significantly shortened as the TNM stage(Pearson’s coefficient = --0.92) and TRPM2 level(Pearson’s coefficient = -0.88) increased in pancreatic cancer patients(Fig. 2c). The progression free survival(PFS) period also significantly shortened as the TNM stage (Pearson’s coefficient = --0.89) and TRPM2 level(Pearson’s coefficient = -0.85) increased in pancreatic cancer patients(Fig. 2c).

The gene set enrichment analysis(GSEA) of human and nude mice PC tissue found that TRPM2 was highly correlated to MEK pathway(Fig. 4d, e, f, Fig. 5a, b, c). According to KEGG database, PKC/MEK was downstream pathway of TRPM2(Fig. 7). The mRNA seq(Fig. 5d), westernblot, and qPCR(Fig. 5e) of 9 different genes in the PKC/MEK pathway found that these genes plus TRPM2 Mwere significantly highly expressed in TT than in TP in PC patients. The BxPC-3 cell model also demonstrated TRPM2 overexpressing was closely related to PKC/MEK pathway in pancreatic cancer(Fig. 6). That was why we concluded TRPM2 could promote PC through PKC/MEK pathway.

MAPK/MEK, which contained molecules like Raf, MEK1/2, and ERK1/2[14], was the downstream pathway of KRAS in PC. KRAS could activate Raf directly, which subsequently promote the development of PC by phosphorylating MEK or ERK in mice[15]. Some researchers tried to block MEK’s activation to inhibit PC development. But AKT/PI3K pathway was accidentally activated after MEK inhibition and resulted in promoting PC^[16]^. If inhibited MEK and PI3K simultaneously, the apoptosis of PC could be increased in vitro assays^[17]^. But this combined inhibition could only work in PC when KRAS was mutated^[18]^. Nowadays, we could find abundant publications about KRAS/MEK’s role in PC. But we still can’t translate them into clinical practice. The exact mechanism still needs to be clarified further.

Protein kinase C(PKC), which also promotes PC development, consists of classical PKC(cPKC), novel PKC(nPKC), and atypical PKC(aPKC)^[19]^. The subfamily was divided by C1 and C2 domain in the molecular structure[20]. cPKC contains both C1 and C2 domain, which could be activated by diacylglycerol(DAG) and Ca^2+^. nPKC contains only C1 domain. It could be activated only by combining DAG. aPKC contains neither C1 nor C2 domain. It could not be activated by DAG or Ca^2+^[20].

PKCα, PKCβ, and PKCγ belongs to cPKC subfamily, in which PKCα Could promote proliferation and metastasis ability of PC by activating Raf-1^[21]^. PKCε, PKCδ, PKCη and PKCθ belongs to nPKC subfamily, in which PKCε and PKCδ could also promote PC by activating Raf-1^[22-24]^. PKCζ and PKCι belongs to aPKC subfamily, in which PKCι could promote PC by activating Rac1-MAPK pathway^[25]^.

TRPM2 is an ion channel on the cell membrane permeable to Na+, K+, and Ca2+. Because TRPM2 is highly expressed in PC, calcium level is also elevated. Calcium could activate PKCα to promote PC further. DAG is an important second messenger in cells. It could be produced in the process of degradation of phospholipids in cell membrane by phospholipase C(PLC)^[26]^. Elevated calcium level in PC could also activate PLC to increase DAG level. Then increased DAG could activate PKCε and PKCδ to promote PC further. PKCι couldn’t be activated by calcium and DAG. So it is not TRPM2 dependent.

## Conclusion

TRPM2 could directly activate PKCα by calcium or indirectly activate PKCε and PKCδ by increased DAG in PC, which promote PC by downstream MAPK/MEK pathway activation(Fig. 8).

**Figure 8.**
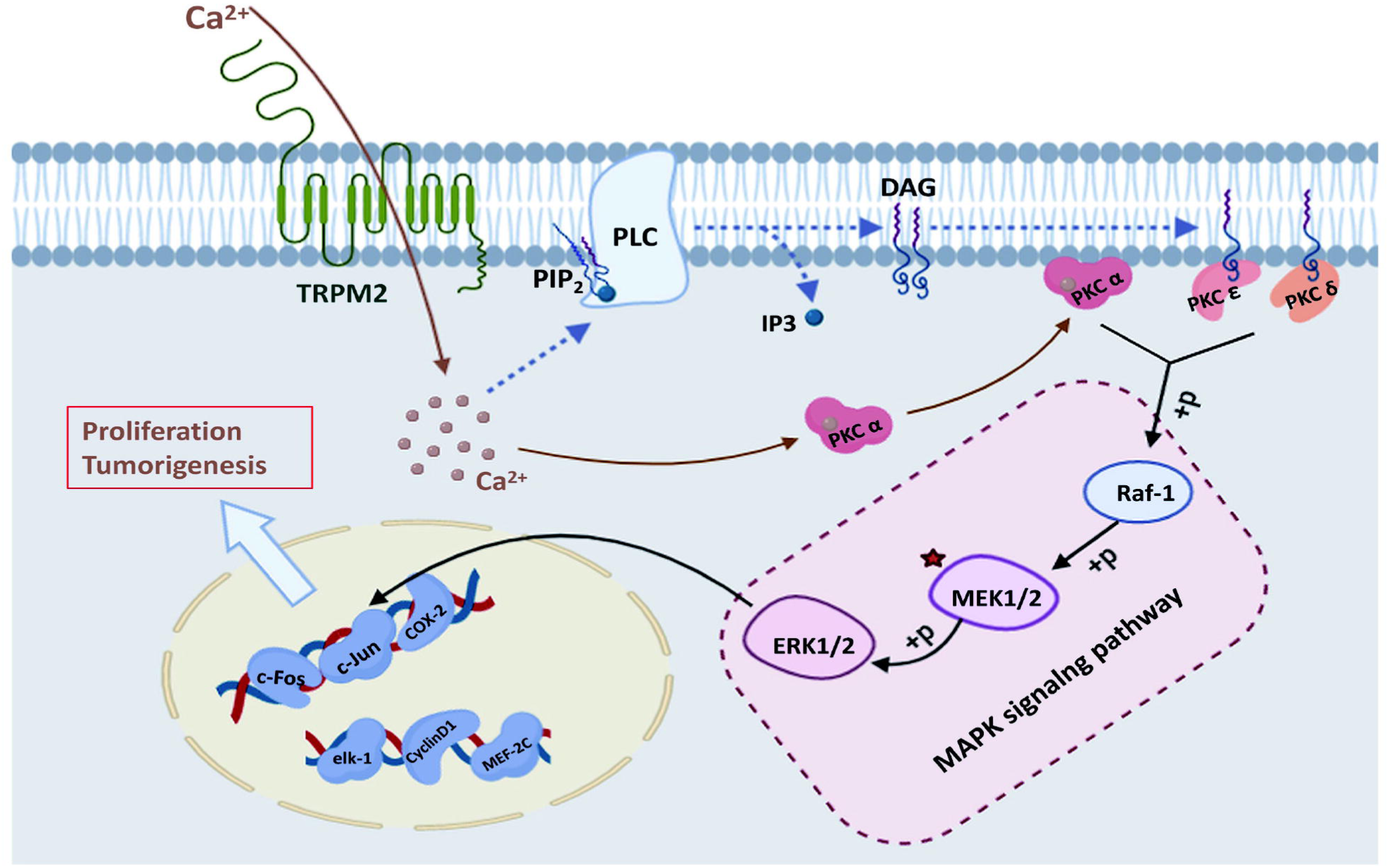
The graph indicates the mechanisms of how TRPM2 promote pancreatic cancer proliferation and tumorigenesis. The increased level of calcium in pancreatic cancer cell could directly activate PKCα, which in turn cause MAPK pathway activation. On the other hand, the increased calcium level could activate MAPK pathway by indirectly activate PKCε and PKCδ.

